# Increased prefrontal activity with aging reflects nonspecific neural responses rather than compensation

**DOI:** 10.1101/156935

**Authors:** Alexa M. Morcom, Richard N. A. Henson, for Cambridge Centre for Ageing and Neuroscience

## Abstract

Elevated prefrontal cortex activity is often observed in healthy older adults despite declines in their memory and other cognitive functions. According to one view, this activity reflects a compensatory functional posterior-to-anterior shift, which contributes to maintenance of cognitive performance when posterior cortical function is impaired. Alternatively, the increased prefrontal activity may be less efficient or less specific, owing to structural and neurochemical changes accompanying aging. These accounts are difficult to distinguish on the basis of average activity levels within brain regions. Instead, we used a novel model-based multivariate analysis technique, applied to two independent functional magnetic resonance imaging datasets from an adult-lifespan human sample (N=123 and N=115; approximately half female). Standard analysis replicated the age-related increase in average prefrontal activation, but multivariate tests revealed that this activity did not carry additional information. The results contradict the hypothesis of a compensatory posterior-to-anterior shift. Instead, they suggest that the increased prefrontal activation reflects reduced efficiency or specificity, rather than compensation.

**Significance statement:** Functional brain imaging studies have often shown increased activity in prefrontal brain regions in older adults. This has been proposed to reflect a compensatory shift to greater reliance on prefrontal cortex, helping to maintain cognitive function. Alternatively, activity may become less specific as people age. This is a key question in the neuroscience of aging. In this study, we used novel tests of how different brain regions contribute to long- and short-term memory. We found increased activity in prefrontal cortex in older adults, but this activity carried less information about memory outcomes than activity in visual regions. These findings are relevant for understanding why cognitive abilities decline with age, suggesting that optimal function depends on successful brain maintenance rather than compensation.

## Introduction

It is well established that healthy aging is associated with a decline in cognitive processes like memory, but mechanistic explanation of this decline is impeded by difficulties in interpreting the underlying brain changes. Functional magnetic resonance imaging (fMRI) of such cognitive processes shows striking increases, as well as decreases, in brain activity of older relative to younger adults. One leading theory – the Posterior-to-Anterior Shift in Aging (PASA) – states that recruitment of anterior regions like prefrontal cortex (PFC) contributes to maintenance of cognitive performance when posterior cortical function is impaired (Davis, Dennis, Daselaar, Fleck, & Cabeza, 2008; Grady, 2012; Park & Reuter-Lorenz, 2009). Alternatively, age-related increases in PFC activity may reflect reduced efficiency or specificity of neuronal responses, reflecting primarily age-related functional impairment within PFC (West, 1996; Glisky et al., 2001; Raz and Rodrigue, 2006; Nyberg, Lövdén, Riklund, Lindenberger, & Bäckman, 2012; Park et al., 2004). It is difficult to adjudicate between these theories based on average activity levels within brain regions (Morcom and Johnson, 2015). We used a novel multivariate approach to directly test predictions of the PASA theory.

With multivariate methods that examine distributed patterns of brain activity over many voxels, one can ask whether increased anterior activity provides additional information, and whether this information goes beyond that provided by posterior cortical regions. Such increases in the information represented by PFC activity would support theories that attribute additional PFC recruitment to compensatory mechanisms. We used a model-based decoding approach called multivariate Bayes (MVB) (Friston et al., 2008; Morcom and Friston, 2012; Chadwick et al., 2014), which estimates the patterns of activity that best predict a target cognitive outcome. Importantly, MVB allows formal comparison of models comprising different brain regions, such as PFC, posterior cortex, or their combination.

In this study, we applied MVB to fMRI data from two different paradigms in population-derived, adult-lifespan samples (N=123 and N=115, 19-88 years; Shafto et al. 2014). In the first, long-term memory (LTM) experiment, participants were scanned while encoding new memories of unique pairings of objects and background scenes, and the target cognitive outcome was whether or not these associations were subsequently remembered (Figure 1a). A previous behavioral study in an independent sample showed a strong decline in such associative memory across the adult lifespan (Henson et al., 2016). To test whether findings generalized across tasks and cognitive domains, as PASA predicts (Davis et al., 2008), we replicated our findings in a second, visual short-term memory (STM) experiment. In the STM experiment, a separate sample of participants was scanned while maintaining visual dot patterns online for a few seconds in the presence of distraction, and the target cognitive outcome was the increase in the number of patterns to be maintained (i.e., load; Figure 1b). Increases in PFC activity have frequently been reported in older adults in similar tasks (Grady et al., 1998; Cabeza et al., 2004), although sometimes activity reductions are found at higher loads (Cappell et al., 2010).

We defined two regions-of-interest (ROIs): posterior visual cortex (PVC), comprising lateral occipital and fusiform cortex, and PFC, comprising ventrolateral, dorsolateral, superior and anterior regions (Fig 2a). These ROIs were based on previous fMRI studies of memory tasks, and those cited in the context of the PASA theory (Davis et al., 2008; Maillet and Rajah, 2014).

**Figure 1.**
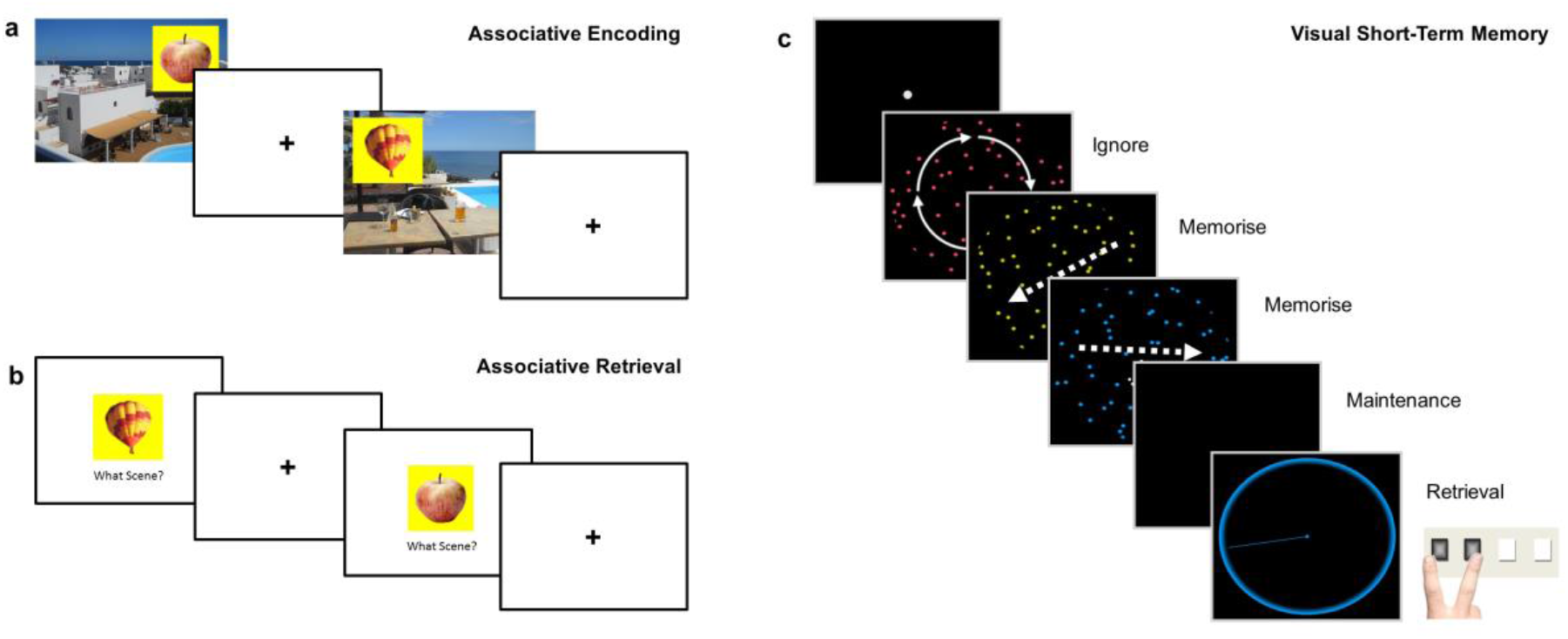
Memory tasks. a and b, long-term memory (LTM) task and c, short-term memory (STM) task. a, Associative encoding. In the scanned Study phase of the LTM task, participants were asked to make up a story that linked each object with its background scene (120 trials total). A scene with positive valence is illustrated. On each trial, the scene was presented for 2 sec, then the object superimposed for 7.5 sec, finally the screen was blanked for 0.5 sec before the next trial. b, Associative retrieval. At Test (out of the scanner), each object was presented again, and after a measure of priming, item memory and background valence memory, participants were asked to verbally describe the scene with which it was paired at Study. The latter verbal recall was scored as correct or incorrect, which was then used to classify the trials at Study into “remembered” and “forgotten” (see text for details). The example illustrates encoding and retrieval of a trial with neutral valence. c, An example trial of STM task with memory load of 2 items. Trials began with fixation dot for 7 sec. On each trial, three dot displays were displayed in red, yellow and blue for 500 msec each (250 msec gap). To vary load, the dots in either one, two or three of the dot displays moved in a consistent direction (the to-be-ignored displays rotated). After the last display, the screen was blanked for an 8 sec maintenance period. Then the probe display presented a colored circle to indicate which dot display to recall (red, yellow or blue). Participants had up to 5 sec to adjust the pointer until the direction matched that of the to-be-remembered display.

## Materials and Methods

### Experiment 1: Long-term Memory Encoding

#### Participants

A healthy, population-derived adult lifespan human sample (N=123; 19-88 years; 66 female) was collected as part of the Cam-CAN study (Shafto et al., 2014). Participants were fluent English speakers in good physical and mental health. Exclusion criteria included a low Mini Mental State Examination (MMSE) score (<=24), serious current medical or psychiatric problems, or poor hearing or vision, as well as standard MRI safety criteria. Two participants were excluded from fMRI analysis as subsequent memory could not successfully be decoded from either region of interest (see Multivariate Bayesian decoding). Two further participants were excluded because of statistical outlier values in the analysis of univariate subsequent memory effects (see Statistics for criteria). The experiment used a within-participant design, so all participants received all the task conditions. Therefore, randomization and blinding were not required. The study was approved by the Cambridgeshire 2 (now East of England— Cambridge Central) Research Ethics Committee. Participants gave informed written consent.

#### Materials

Stimuli were 160 pictures of everyday emotionally-neutral objects taken from Smith et al. (2004). For the study phase, objects were presented within a square yellow background on one of 120 scenes from the IAPS emotional pictures database (Lang et al., 1997). Scenes were grouped into 40 per valence (positive, neutral, negative), selected based on a pilot study, with the same randomized trial order for each valence condition for all participants. To control for stimulus effects, the 160 objects were divided randomly into 4 sets, and the allocation of object sets to scene valence rotated across participants in 4 different counterbalances (see Henson et al., 2016, for further details).

#### Behavioral procedure

The paradigm is summarized in Figure 1a. The scanned study phase comprised 120 trials, presented in two 10 min blocks separated by a short break. On each study trial, a background scene was first presented for 2 sec, and an object then superimposed for 7.5 sec, slightly above center and either to the left or right. Participants were asked to create a story that linked the object to the scene, to press a button when they had made the story. In order to equate the amount of time spent processing the story, participants were asked to continue to elaborate it until the scene and object disappeared. A blank screen of 0.5 sec was then presented prior to the next trial. Participants were informed that the task would include some pleasant and unpleasant scenes. They were not told that their memory would be tested later. A practice session of 6 study trials was given just beforehand.

The test phase took place outside the scanner, following a short break of approximately 10 min involving refreshment and conversation with the experimenter. The 120 objects from the study phase were presented again, randomly intermixed with 40 new objects, and divided into 4 blocks lasting approximately 20 min each. The first stage of each test trial involved tests of priming, item memory and memory for the picture valence (see Henson et al., 2016 for details) However, for the present fMRI analysis, we focused on the fourth question in each trial. Participants were asked to verbally recall the scene that had been paired with the test object at study. Trials at study in which scenes that were correctly recalled at test, in terms of detail or gist, were scored as “remembered”. Remaining trials were split according to whether the scenes could not be recalled (“associative misses”), or for which an incorrect scene was described instead (“associative intrusions”), or for which the object was not recognized (“item misses”). Initial analyses showed no evidence that valence interacted with age, so trials were collapsed across valence (see Behavioral results). Table 1 summarizes the trial numbers per condition split by age tertile.

**Table 1.** Trial numbers divided by condition. Remembered = trials with correct object recognition and scene recall; Associative Miss = trials with correct object recognition but no scene recall; Associative Intrusion = trials with correct object recognition but recall of an incorrect scene; Item Miss = trials with misclassification of the object as unstudied. Data are split by age tertile. Means are given with standard deviations (SD).

|  | Younger<br>(19-45 years)<br>n=38 | Middle-Age<br>(46-64 years)<br>n=43 | Older<br>(65-88 years)<br>n=42 |
| --- | --- | --- | --- |
| Remembered | 55<br>(25) | 44<br>(24) | 23<br>(15) |
| Forgotten |  |  |  |
| Associative Miss | 31<br>(13) | 38<br>(17) | 46<br>(18) |
| Associative Intrusion | 11<br>(8) | 13<br>(10) | 10<br>(8) |
| Item Miss | 22<br>(15) | 25<br>(17) | 42<br>(24) |

For the main imaging analyses, we combined the three types of forgotten trial in order to maximise power. However, in case processes that lead to subsequent memory for associative memory versus item memory differ (e.g., Dennis et al., 2008; Mattson et al., 2014), we ran a subsidiary imaging analyses with item misses excluded.

#### Imaging data acquisition and preprocessing

The MRI data were collected using a Siemens 3 T TIM TRIO system (Siemens, Erlangen, Germany). MR data preprocessing and univariate analysis used the SPM12 software (Wellcome Department of Imaging Neuroscience, London, UK, www.fil.ion.ucl.ac.uk/spm), release 4537, implemented in the AA 4.0 pipeline (https://github.com/rhodricusack/automaticanalysis). The functional images were acquired using T2*-weighted data from a Gradient-Echo Echo-Planar Imaging (EPI) sequence. A total of 320 volumes were acquired in each of the 2 Study sessions, each containing 32 axial slices (acquired in descending order), slice thickness of 3.7 mm with an interslice gap of 20% (for whole brain coverage including cerebellum; TR =1970 msec; TE = 30 msec; flip angle =78 degrees; field of view (FOV) =192 mm × 192 mm; voxel-size = 3 mm × 3 mm × 4.44 mm). A structural image was also acquired with a T1-weighted 3D Magnetization Prepared RApid Gradient Echo (MPRAGE) sequence (repetition time (TR) 2250ms, echo time (TE) 2.98 ms, inversion time (TI) 900 msec, 190 Hz per pixel; flip angle 9 deg; FOV 256 x 240 x 192 mm; GRAPPA acceleration factor 2).

The structural images were rigid-body registered with an MNI template brain, bias-corrected, segmented and warped to match a gray-matter template created from the whole CamCAN Stage 3 sample (N=272) using DARTEL (Ashburner, 2007) (see Taylor et al., 2015) for more details). This template was subsequently affine-transformed to standard Montreal Neurological Institute (MNI) space. The functional images were then spatially realigned, interpolated in time to correct for the different slice acquisition times, rigid-body coregistered to the structural image and then transformed to MNI space using the warps and affine transforms from the structural image, and resliced to 3x3x3mm voxels.

#### Univariate imaging analysis

For each participant, a General Linear Model (GLM) was constructed, comprising three neural components per trial: 1) a delta function at onset of the background scene, 2) an epoch of 7.5 sec which onset with the appearance of the object (2 sec after onset of scene) and offset when both object and scene disappeared, and 3) a delta function for each keypress. Each neural component was convolved with a canonical haemodynamic response function (HRF) to create a regressor in the GLM. The scene onset events were split into 3 types (i.e, 3 regressors) according to the valence of the scene on each trial, while the keypress events were modelled by the same regressor for all trials (together, these four regressors served to model trial-locked responses that were not of interest). The responses of interest were captured by the epoch neural component, during which participants were actively relating the scene and object (see Behavioral Procedure). The duration of this component did not depend on response time, as participants were instructed to continue to link the object and scene mentally for the full duration of the display.

For the principal GLMs, the epoch component was split into 6 types (regressors) according to the 3 scene valences and 2 types of subsequent memory, i.e., study trials for which the scenes were correctly recalled (“remembered”), and those for which the scenes could not be recalled, an incorrect scene was described instead, or the object was not recognized (“forgotten”). When comparing remembered and forgotten trials, we averaged across the three valences because 1) there was no behavioral evidence of an interaction between age and valence on subsequent memory, 2) there was no imaging evidence of an interaction between age and valence on subsequent memory, and 3) there would have been insufficient numbers of each trial-type to examine each valence separately. Thus the main target contrast for the univariate and multivariate analyses were remembered versus forgotten trials.

As noted above, given that encoding of associative (source) information versus item information may differ with regard to additional recruitment and (potentially) to compensation (e.g., Dennis et al., 2008; Mattson et al., 2014), we ran a subsidiary analysis in which the “forgotten” category excluded item misses. In these GLMs, study trials were modelled using 9 regressors according to the 3 scene valences and 3 (rather than 2) types of subsequent memory: trials on which the object was recognized but the scene forgotten or incorrectly recalled (“association forgotten”) and trials on which the object was not recognized (“item forgotten”). Participants for whom one of the sessions did not contain at least one trial of each type were removed, leaving n=109 (note this involved removal of more participants in the oldest age tertile: 0 removed aged 19-35, 2 aged 46-64, and 12 aged 65-88 years). One remaining outlier (>5 SD) on the univariate measures was removed, giving n=108. As reported in the Results section, this subsidiary analysis corroborated the findings of the main analysis.

Six additional regressors representing the 3 rigid body translations and rotations estimated in the realignment stage were included in each GLM to capture residual movement-related artifacts. Finally the data were scaled to a grand mean of 100 over all voxels and scans within a session, and the model was fitted to the data in each voxel. The autocorrelation of the error was estimated using an AR(1)-plus-white-noise model, together with a set of cosines that functioned to highpass the model and data to 1/128 Hz, fit using Restricted Maximum Likelihood (ReML). The estimated error autocorrelation was then used to “prewhiten” the model and data, and ordinary least squares used to estimate the model parameters. To compute subsequent memory effects, the parameter estimates for the 6 epoch components were averaged across the two sessions and the three valences (weighted by number of trials per session/valence), and contrasted directly as remembered minus forgotten (Morcom et al., 2003; Maillet and Rajah, 2014). Univariate statistical analyses were conducted on the mean subsequent memory effect across all voxels in the MVB analysis, in each ROI for each participant (see next section).

### Experiment 2: Visual STM

#### Participants

Participants were a separate subset (N=115; 25-86 years; 54 female) of those recruited to the Cam-CAN study (see Experiment 1, Participants, for details). Nineteen participants were excluded from the current analysis as the contrast of interest could not successfully be decoded from either region of interest (see Multivariate Bayesian decoding). None were excluded because of statistical outlier values on the measures used (see Statistics for criteria). The experiment used a within-participant design, so all participants received all the task conditions.

#### Materials

The task was adapted from Emrich et al. (2013). Stimuli were three patches of coloured dots, one red, one yellow, and one blue. Dots were 0.26 degrees of visual angle (dva) in diameter, at a density of 0.7 per square degree, and viewed though a circular aperture of diameter 11 dva. As a manipulation of set size, one, two, or three of the dot displays moved (at 2 dva/ sec) in a single direction which had to be remembered. The other, distractor, displays rotated around a central axis, and were be ignored. On 90% of trials the probed movements were in one of three directions (7, 127, or 247 degrees). Other directions were selected at random, to avoid subjects learning the target directions. Order of presentation of the 3 display colors was randomized trial by trial, as was memory load. Rotation direction alternated across trials of a given load.

#### Behavioral procedure

Each trial began with a grey fixation dot in the middle of a black screen for 5 sec, which then turned white for 2 sec. Participants then saw the 3 dot displays for 500 msec each, with 250 msec in-between. After the third display, an 8 sec blank fixation delay was presented, followed by the probe display. The probe display showed a colored circle to indicate which dot display to recall (red, yellow, or blue), with a pointer. Participants had up to 5 sec to adjust the pointer using 2 buttons until it matched the direction of motion of the remembered target dot display.

After responding, a third button cleared the probe display. Participants completed 3 runs of 30 trials per run (10 for each load). The direction of the target, the sequential position of the target, and the set size were counterbalanced within each run, and presented in random order. Colour and position of target were also counterbalanced using a Greco-Latin square design.

#### Imaging data acquisition and preprocessing

Imaging data were acquired on the same scanner as Experiment 1. Functional T2*-weighted data were acquired using a Multi-Echo Gradient-Echo Echo-Planar Imaging (EPI) sequence. Approximately 300 volumes were acquired in each of the 3 VSTM task sessions (duration depending on response times). Each volume had 34 axial slices (acquired in descending order), slice thickness of 2.9 mm with an interslice gap of 20% (FOV = 224 mm × 224 mm, TR = 2000 msec; TE = 12 msec, 25 msec, 38 msec; flip angle = 78 degrees; voxel-size = 3.5 mm × 3.5 mm × 3.48 mm). Structural image sequences were the same as in Experiment 1. The multiple echos were combined by computing their average, weighted by their estimated T2* contrast. The functional images were spatially realigned and interpolated in time to correct for different slice acquisition times. Spatial normalisation used the ‘new segment’ protocol in SPM12 (Ashburner and Friston, 2005). Participants’ structural scans were coregistered to their mean functional image, then segmented into 6 tissue classes. Functional images were rigid-body coregistered to the structural image then transformed to MNI space using the warps and affine transforms estimated from the structural image, and resliced to 2x2x2mm voxels.

#### Univariate imaging analysis

The GLM for each participant comprised three neural components per trial: 1) encoding, modelled as an epoch of 1 sec duration at onset of the first moving dot pattern, 2) maintenance, modelled as an epoch of 4 sec at offset of the last moving dot pattern (2.25 sec after onset of 1), and 3) probe, a delta function at the time of the participant‘s response. These components were each split into 3 types (regressors) according to the 3 STM load levels. As in Experiment 1, 6 additional regressors were added representing the motion parameters estimated in the realignment stage. Finally the data were scaled to a grand mean of 100 over all voxels and scans within a session. To confirm that this dataset was suitable as a replication of Experiment 1’s multivariate results, we first checked that at least one significant cluster within the PFC region of interest (ROI) showed increased univariate activity in older people. This was done using a standard analysis of effects of a linear contrast of increasing VSTM load on activity during the delay period, whole-brain corrected for multiple comparisons at p < .05 (voxel threshold), and a linear contrast of age. Details of this analysis are not reported and it did not contribute to ROI selection, which was the same as for Experiment 1. Note that the continuous nature of the judgment in the VSTM task precludes definition of individual trials as correct or incorrect (rather, performance is used to estimate continuous summary measures for each participant, as in Emrich et al, 2013). Therefore all trials were included in the fMRI analysis, and the main target contrast for the univariate and multivariate analyses was the linear effect of load from 1-3 during the delay period.

### Regions of interest

ROIs were defined using WFU PickAtlas (http://fmri.wfubmc.edu/, version 3.0.5) with AAL and Talairach atlases (Lancaster et al., 2000; Tzourio-Mazoyer et al., 2002; Maldjian et al., 2003). The posterior visual cortex (PVC) mask comprised bilateral lateral occipital cortex and fusiform cortex (from AAL, fusiform and middle occipital gyri), and the PFC mask comprised bilateral ventrolateral, dorsolateral, superior and anterior regions: from AAL, the inferior frontal gyrus (IFG), both pars triangularis and pars orbitalis; middle frontal gyrus, lateral part (MFG); superior frontal gyrus (SFG); and from Talairach, Brodmann Area 10 (BA10), dilation factor = 1. In the subregion analyses, separate masks were created for BA10, IFG, MFG and SFG (regions included in the BA10 mask were excluded from the other masks).

### Multivariate Bayesian decoding

A series of MVB decoding models were fit to assess the information about subsequent memory carried by individual ROIs or combinations of ROIs. Each MVB decoding model is based on the same design matrix of experimental variables used in the univariate GLM, but the mapping is reversed: many physiological data features (derived from fMRI activity in multiple voxels) are used to predict a psychological target variable (Friston et al., 2008). This target (outcome) variable is specified as a contrast. In Experiment 1 (LTM) the outcome was subsequent memory, and in Experiment 2 (STM) it was the linear increase in STM load during maintenance periods. Modelled confounds in the design (all covariates apart from those involved in the target contrast) are removed from both target and predictor variables.

Each MVB model is fit using hierarchal parametric empirical Bayes, specifying empirical priors on the data features (voxel-wise activity) in terms of patterns over voxel features and the variances of the pattern weights. Since decoding models operating on multiple voxels (relative to scans) are ill-posed, these spatial priors on the patterns of voxel weights act as constraints in the second level of the hierarchical model. MVB also uses an overall sparsity (hyper) prior in pattern space which embodies the expectation that a few patterns make a substantial contribution to the decoding and most make a small contribution. The pattern weights specifying the mapping of data features to the target variable are optimised with a greedy search algorithm using a standard variational scheme which iterates until the optimum set size is reached (Friston et al., 2007). This is done by maximizing the free energy, which provides an upper bound on the Bayesian log-evidence (the marginal probability of the data given that model). The evidence for different models predicting the same psychological variable can then be compared by computing the difference in their log-evidences, giving the log (marginal) likelihood ratio test (Bayes factor) (see Friston et al., 2007; Chadwick et al., 2012; Morcom and Friston, 2012). In this work, the main outcome measures were the log-evidence for each model and the set of fitted weights for all patterns (voxels) in the ROI, which can be examined to assess their distribution over voxels and the contributions of different combinations of voxels. These analyses were implemented in SPM12 v6486 and custom MATLAB scripts.

Features (voxels) for MVB analysis were selected using an orthogonal contrast and a leave-one-participant-out scheme. For each participant and ROI, these were the 1000 voxels with the strongest responses to the task: in Experiment 1 (LTM), the 6 epoch regressors modelling object onsets in the GLM, and in Experiment 2 (STM), the 3 epoch regressors modelling maintenance periods in the GLM (defined using an F contrast in all other participants testing variance explained by these regressors, regardless of valence or subsequent memory). We used a sparse spatial prior, in which each pattern is an individual voxel (Morcom and Friston, 2012; Chadwick et al., 2014; Hulme et al., 2014; Maass et al., 2014). We first checked that the target memory variables could reliably be decoded from the selected features by contrasting the evidence for each model with the evidence for models in which the design matrix (and therefore the target variable) had been randomly phase-shuffled, taking the mean over 20 repetitions, and comparing log-evidence for real versus phase-shuffled models. One-tailed t-tests compared the difference in real versus shuffled model evidences to a hypothesized population mean difference of 3 which would reflect good Bayesian evidence for the real over the shuffled models. These showed that the difference in log-evidence was robustly greater than this in both PVC, *t*(118) = 6.08, p < .0001, *r*^2^_adj_ = .225, mean difference = 9.72; and PFC, *t*(118) = 7.70, p < .0001, *r*^2^_adj_ = .323, mean difference = 11.8. The same applied to Experiment 2: for PVC, *t*(95) = 8.42, p < .0001, *r*^2^_adj_ = .415, mean difference = 23.0, and PFC, *t*(95) = 11.4, p < .0001, *r*^2^_adj_ = .569, mean difference = 35.0. To confirm that the sparse prior represented the best spatial model, we then compared the log-evidence with that for models with smooth spatial priors, in which each pattern is a local weighted mean of voxels (Gaussian FWHM = 8). For Experiment 1 (LTM): log evidence was substantially greater for the sparse priors in both ROIs: in PVC, *t*(118) = 18.4, p < .0001, *r*^2^_adj_ = .737, and PFC, *t*(118) = 18.0, *r*^2^_adj_ = .728, p < .0001, two-tailed tests. The same was true for Experiment 2 (STM): for PVC, *t*(95) = 10.3, *r*^2^_adj_ = .464, p < .0001, and PFC, *t*(95) = 14.9, *r*^2^_adj_ = .650, p < .0001.

Unlike univariate activation measures such as subsequent memory effects, but like other pattern-information methods, MVB finds the best non-directional model of activity predicting the target variable, so positive and negative pattern weights are equally important. Therefore, the principle MVB measure of interest for each ROI was the spread (standard deviation) of the weights over voxels, reflecting the degree to which multiple voxels carried substantial information about subsequent memory. To test whether PFC activity was compensatory, we also constructed a novel measure of the contribution of prefrontal cortex to subsequent memory. This used Bayesian model comparison within participants to assess whether a joint PVC-PFC model boosted prediction of subsequent memory relative to a PVC-only model. The PASA hypothesis, in which PFC is engaged to a greater degree in older age and this contributes to cognitive outcomes, predicts that a boost will be more often observed with increasing age. The initial dependent measure was the log model evidence, coded categorically for each participant to indicate the outcome of the model comparison. The 3 possible outcomes were: a boost to model evidence for PVC-PFC relative to PVC models, i.e., better prediction of subsequent memory (difference in log evidence > 3), equivalent evidence for the two models (-3 < difference in log evidence < 3), or a reduction in prediction of subsequent memory for PFC-PVC relative to PVC (difference in log evidence < -3).

Lastly, given that the relative contribution of anterior versus posterior regions could change with age, even if the absolute amount of pattern information decreased with age in both regions, we computed a second novel measure: we estimated the PFC contribution to cognitive outcome in terms of the proportion of top-weighted voxels in the joint PVC-PFC model that were located in PFC, as opposed to PVC, derived from joint PVC-PFC models. In each participant, the voxels making the strongest contribution to the cognitive outcome, defined as those with absolute voxel weight values greater than 2 standard deviations from the mean, were split according to their anterior versus posterior location. The dependent measure was the proportion of these top voxels located in PFC.

### Experimental design and statistical analysis

Sample size was determined by the initial considerations of Stage 3 of the CamCAN study – see Shafto et al.(Shafto et al., 2014) for details. For the LTM experiment, a sensitivity analysis indicated that with N=123, we would have 80% power to detect a small to medium effect explaining 6.5% of the variance on a two tailed test for a model with 2 predictors (*r*^2^ = .0658). For the STM experiment with N = 115, the corresponding minimal effect size for 80% power was 6.9% of the variance (*r*^2^ = .0694). In our previous report of aging and successful memory encoding, an a *priori* test of a between-region difference in subsequent memory effects according to age showed a large effect (*r*^2^ = .257) (Morcom et al., 2003).

Age effects on continuous multivariate or univariate dependent measures were tested using robust second-order polynomial regression with “rlm” in the package MASS for R (Venables et al., 2002); MASS version 7.3-45; R version 3.3.1) with standardized linear and quadratic age predictors. For analysis of covariance for behavioral data we used JASP version 0.8.3.1. Analysis of outcomes of the between-region MVB model comparison (PVC and PFC combined versus PVC, see Fig 2 and main text) used ordinal regression with “polr” in MASS. Distributions were also trimmed to remove extreme outliers (> 5 SD above or below the mean). In Experiment 1 (LTM), the two participants (aged 72 and 80) with outlier values for univariate effects were also removed from the MVB analyses so the samples examined were comparable. We excluded two further participants (aged 68 and 83) in whom subsequent memory could not be decoded from at least one of the two ROIs (log model evidence <= 3), giving n=119. In Experiment 2 (STM), we excluded nineteen participants in whom VSTM load could not be decoded during maintenance, giving n=96. All tests were two-tailed and used an alpha level of .05.

Where it was important to test for evidence for the null hypothesis over an alternative hypothesis, we supplemented null-hypothesis significance tests with Bayes Factors (Wagenmakers, 2007; Rouder et al., 2009). The Bayes Factors were estimated using Dienes’ online calculator (Dienes, 2014) which operationalizes directional hypotheses such as PASA in terms of a half-normal distribution. Here, we assumed an effect size of 1 SD and therefore defined the half-normal distribution with mean=0 and SD=1. All statistics and p values are reported to 3 significant figures, except where p < .0001.

## Results

### Experiment 1: Long-term Memory (LTM) Encoding

#### Behavioral results

We examined age effects on the number of trials in each remembered and forgotten condition (see Table 2). For remembered trials, there was a significant linear decrease with age (*t*(118) = -7.30, p < .0001, *r*^2^_adj_ = .299), with no significant quadratic component (*t*(118) = -0.104, p = .917; alpha = .0125). As a consequence, the number of forgotten trials increased with age, and this was true for both associative misses (linear *t*(118) = 4.82, p < .0001, *r*^2^_adj_ = .150; quadratic, *t*(118) = 0.630, p = .532) and item misses (linear *t*(118) = 5.43, p < .0001, *r*^2^_adj_ = .186; quadratic, *t*(118) = 1.57, p = .120), although not for associative intrusions (linear *t*(118) = -1.38, p = .163; quadratic, *t*(118) = -2.29, p = .0221). Analysis of covariance with the factor of valence (Positive, Neutral, Negative) showed no interaction between age and valence on the number of remembered items (for linear effect of age, *F*(2,231) < 1, p = .486; quadratic, *F*(2,231) = 1.59, p = .206).

#### Testing compensation

Standard univariate activation analyses assessed mean activity in each ROI across all voxels included in the multivariate analysis (see Materials and Methods). Consistent with the PASA account, the increase in activity associated with subsequent memory became more pronounced with age, particularly in later years (linear effect of age, *t*(118) = 2.43, p = .0166; quadratic effect of age, *t*(118) = 2.58, p = .0111) (Fig 2a, Table 2). Age effects in PVC were not significant (see Table 2). The age effects in PFC were also present after removal of the older participant with the largest SM effect (although they did not meet our criterion for an outlier; see Fig 2a; linear *t*(117) = 2.14, p = ; quadratic *F*(117) = 2.31, p = .033). In both ROIs results were very similar for the models excluding forgotten trials for which the items themselves were forgotten (see Methods; PFC: linear *t*(107) = 2.22, p = .0316; quadratic *t*(107) = 2.91, p = .00527; PVC: linear, *t*(107) = 1.10, p= .288; quadratic, *t*(107) = 1.24, p = .233).

**Figure 2.**
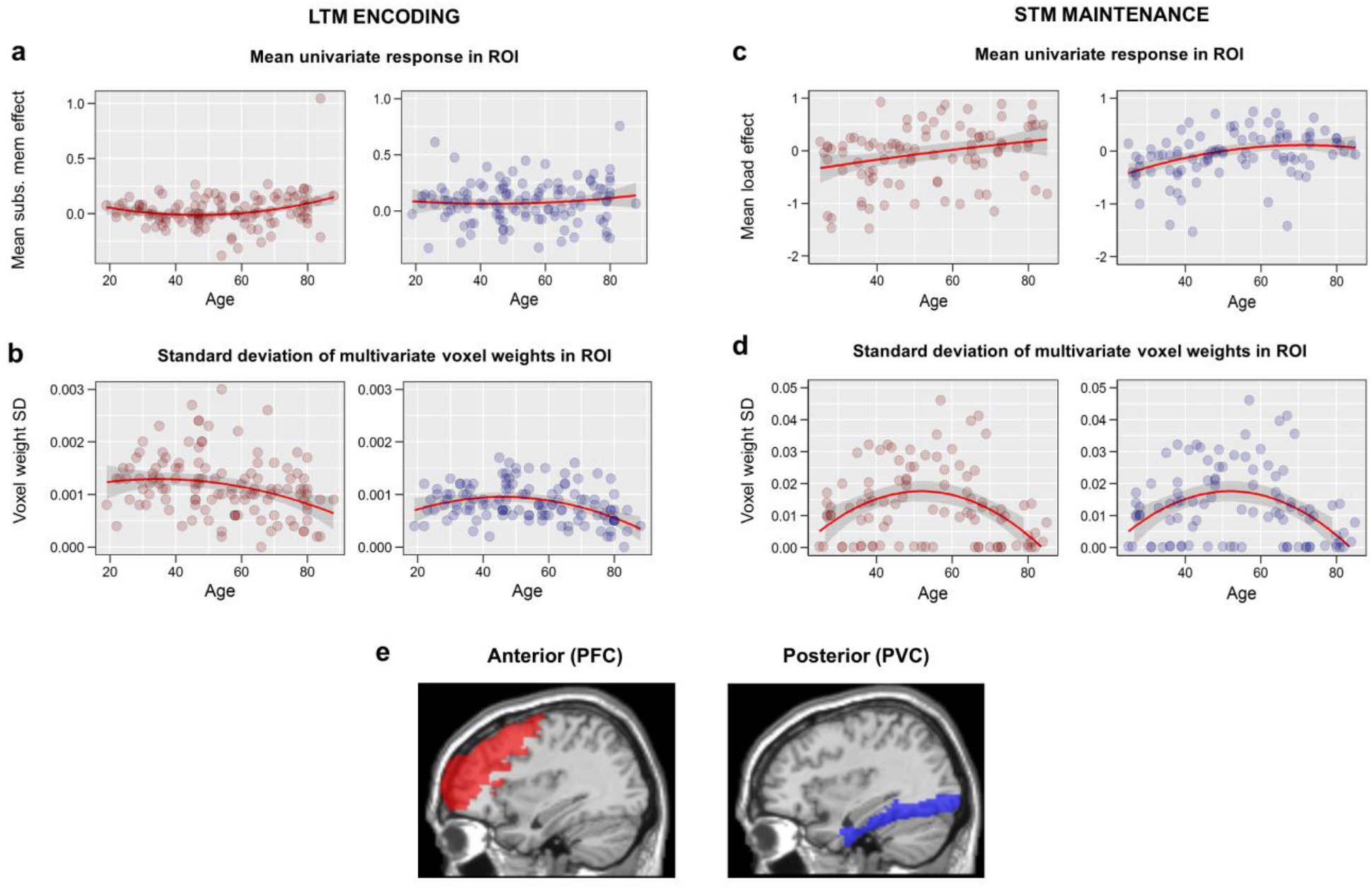
Relationship between age and univariate and multivariate effects within ROIs. **(a)**. Univariate subsequent memory effects (mean activity for remembered - forgotten), showing increased activity with age in PFC but not PVC. **(b)**. Spread of multivariate responses predicting subsequent memory (standard deviation of fitted MVB voxel weights), showing reduced spread of responses with age in both ROIs. **(c)**. Univariate effects of load (positive linear contrast) during STM maintenance, showing increased activity with age in both ROIs. **(e)**. Spread of multivariate responses during STM maintenance predicting increasing load, showing reduced spread of responses with age in both ROIs. Red and blue lines are robust-fitted second-order polynomial regression lines and shaded areas show 95% confidence intervals. **(e)**. PVC (blue) and PFC (red) ROIs overlaid on sagittal section (x=+36) of a canonical T1 weighted brain image. Note that y-axis scales are not comparable across tasks.

**Table 2.** Age effects on mean univariate SM effects and spread of multivariate SM effects in the LTM task. PFC-PVC refers to analyses where the dependent variable was the difference in each measure between PFC and PVC. SD = standard deviation. *r*^2^_adj_ = the unbiased estimate of the amount of variance explained in the population. n = 119.

| ROI/<br>measure | Model |  | Linear |  |  | Quadratic |  |  |
| --- | --- | --- | --- | --- | --- | --- | --- | --- |
| | *F* | *p* | *t* | $r^2_{adj}$ | *p* | *T* | $r^2_{adj}$ | *p* |
| Mean univariate SM activation |  |  |  |  |  |  |  |  |
| PFC | 5.49 | .00525 | 2.43 | .0312 | 0.0166 | 2.58 | .0371 | .0111 |
| PVC | 0.426 | .654 | 0.728 | -- | 0.480 | 0.703 | -- | .495 |
| PFC-PVC | 0.837 | .436 | 0.883 | -- | 0.388 | 1.084 | -- | 0.293 |
| Multivariate spread (SD) of SM activity |  |  |  |  |  |  |  |  |
| PFC | 6.36 | .00240 | -3.33 | .0701 | .000998 | -1.44 | -- | .151 |
| PVC | 11.3 | < .0001 | -3.49 | .0780 | .000650 | -3.50 | .0784 | .000621 |
| PFC-PVC | 2.02 | .109 | 4.16 | -- | .0437 | .398 | -- | .690 |

If the increasing PFC activation with age reflected compensation, we expected the multivariate analyses to show that this increased activity carried additional information about subsequent memory. However, the data revealed a different pattern. In MVB models, like other linear models with multiple predictors, each voxel within an ROI has a weight that captures the unique information it contributes (in this case, for predicting subsequent memory). Because both positive and negative weights carry information, we summarised the MVB results by the spread (standard deviation) of weights over voxels (see Materials and Methods).

In both ROIs, this spread was markedly reduced during later life (PVC: linear *t*(118) = -3.49, p= .000650; quadratic *t*(118) = -3.50, p = .000621; PFC: linear *t*(118) = -3.33, p = .000998; quadratic *t*(118) = -1.44, p = .151); see Fig 2b and Table 2. This means that, contrary to a compensatory PASA shift, PFC showed fewer, rather than more, voxels with large positive or negative weights with increasing age. Again, the results were similar for the subsidiary models excluding item misses (PVC: linear *t*(107) = -1.41, p= .158; quadratic *t*(107) = -2.81, p = .000544; PFC: linear *t*(107) = -2.21, p = .0280; quadratic *t*(107) = -0.566, p = .570). By contrast, the spread of univariate activities across voxels increased with age in both ROIs (for PVC, linear effect of age, *t*(118) = 5.91, p < .0001, *r*^2^_adj_ = .215; quadratic effect of age, *t*(118) = 1.72, p = .0881; for PFC, linear effect of age, *t*(118) = 5.64, p < .0001, *r*^2^_adj_ = .199; quadratic *t*(118) = -2.31, p = .0226, *r*^2^_adj_ = .0268).

To provide a more direct test for a compensatory posterior-to-anterior shift, we assessed the specific contribution of PFC to subsequent memory, over and above that of PVC. We fitted a joint MVB model that included both posterior and anterior ROIs, and contrasted this with a model including PVC only, using Bayesian model comparison. Thus we could ask, for each participant, whether or not adding PFC to the model “boosted” prediction of subsequent memory (see Methods). Contrary to the PASA theory of a compensatory shift towards greater reliance on PFC, a Bayes Factor comparing these two models revealed strong evidence for the null hypothesis of no boost (BF_01_ = 11.1); indeed, the probability of a boost to model evidence for PVC-PFC compared to PVC-only actually decreased with age numerically (Fig 3a; linear *t*(118) = -1.54, p = .126). Excluding item misses from the model enhanced this pattern (for linear age effect *t*(107) = -2.34, p = .0211, BF_01_ = 14.3).

**Figure 3.**
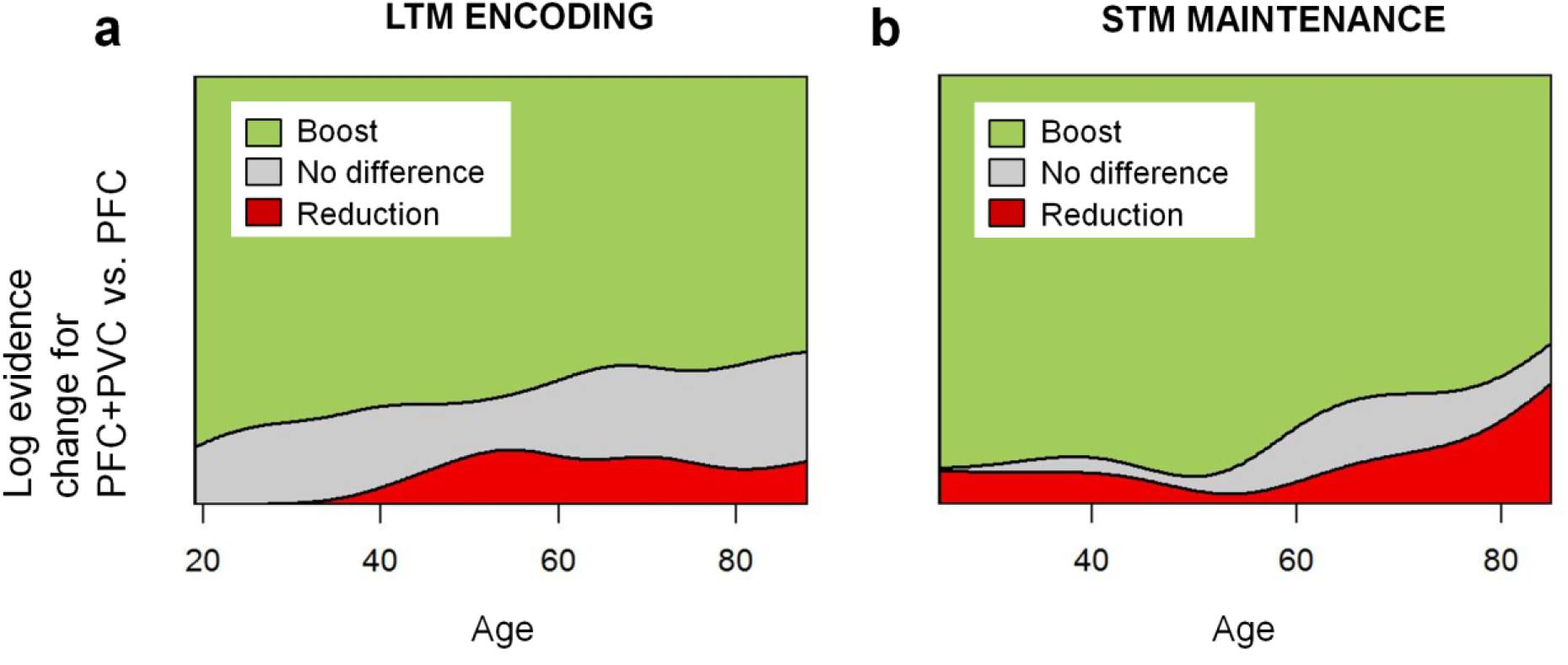
Evidence against a compensatory posterior-to-anterior shift from MVB comparisons between ROIs. Ordinal regression of Bayesian model comparison of combined PVC+PFC model versus PVC-only model using age to predict outcomes of model comparison: adding PFC to the model boosts prediction of the cognitive outcome (difference in log-evidence > 3), or there is no boost (-3 < difference < 3), or a reduction in log-evidence (difference < -3). (a) LTM, for subsequent memory effects, a boost was no more frequent with increasing age (b) STM, for load effects, a boost was less frequent with increasing age.

#### Testing sub-regions within PFC

We also explored whether the pattern of results was similar across subregions within PFC. The PASA theory does not specify which areas are involved in a compensatory shift, but aging does not affect all subregions equally (Raz and Rodrigue, 2006). In functional studies, univariate SM effects in ventrolateral and dorsolateral PFC tend to be age-invariant while anterior and superior prefrontal regions show age-related increases (Morcom et al., 2003; Maillet and Rajah, 2014). In contrast, Davis et al.’s (2008) PASA proposal was based on increased activation in older people in anterior ventrolateral PFC and anterior cingulate during visual perception and episodic retrieval. In the present episodic encoding task, there were significant age-related increases in univariate activation in anterior PFC (BA10) and lateral inferior frontal gyrus (IFG), and significant decreases in the spread of multivariate voxel weights in BA10 and superior (medial) frontal gyrus (SFG), as well as numerical decreases also in IFG and in lateral middle frontal gyrus including dorsolateral PFC (MFG) (Table 3; see Materials and Methods, Regions of Interest for region definition). Thus the overall age-related increase in activation was mainly driven by BA10 and IFG, but no subregion showed a decrease in activation with age. The reduction in multivariate information and evidence against a functional boost were relatively uniform over subregions (Table 3). Direct model comparison showed no evidence that PFC activity was compensatory in older age, even in the two subregions with strong increases in activation: Bayes Factors weighed against any boost to prediction of subsequent memory for joint PFC-PVC models relative to PVC-only models (BF_01_ favoring the null over positive linear effect of age = 11.1 for BA10, 12.5 for MFG, 5.00 for IFG and 14.2 for SFG).

**Table 3.**
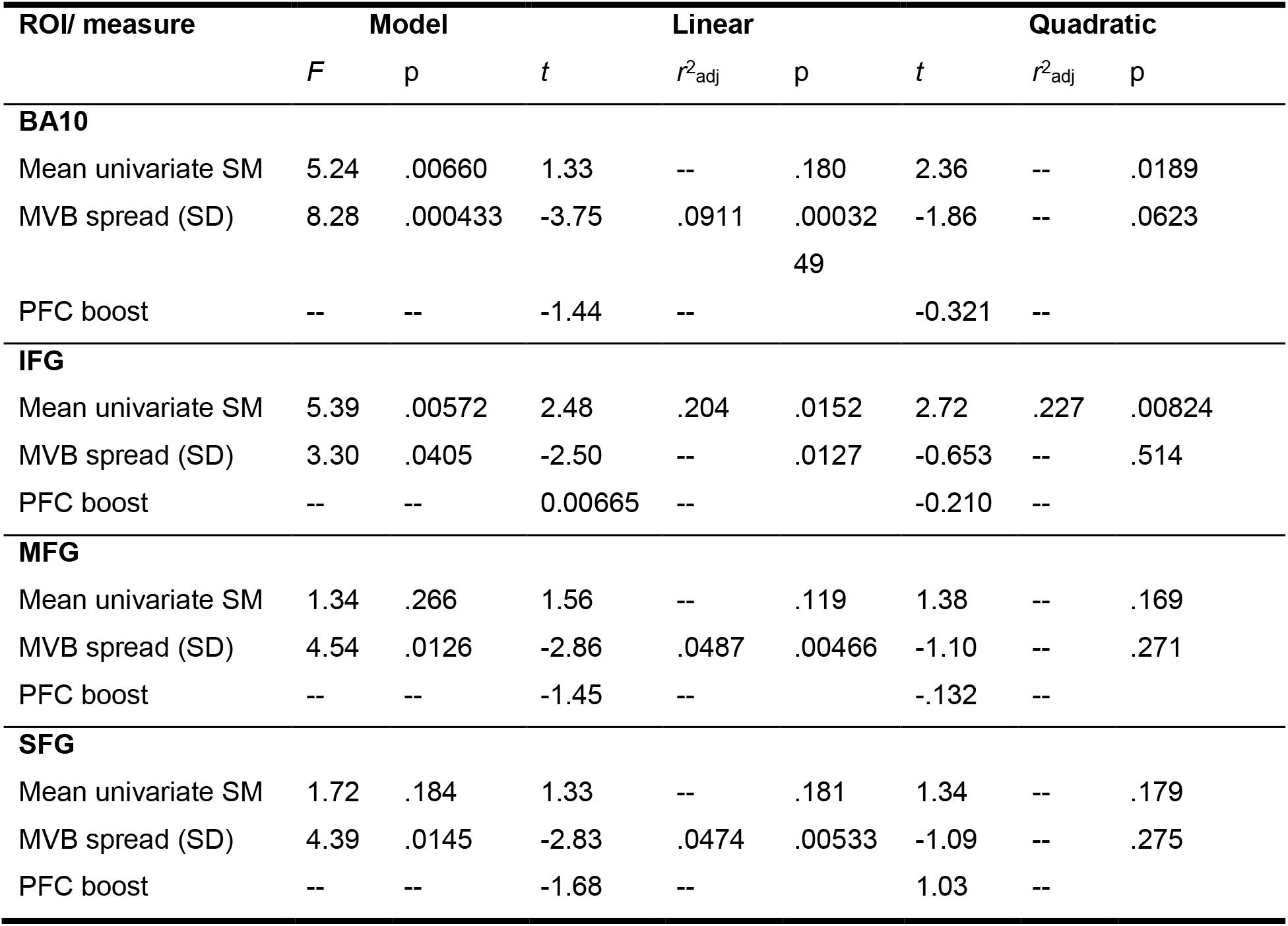
Age effects for PFC subregions in the LTM task. The table lists mean univariate SM effects, the spread (SD) of multivariate Bayesian (MVB) voxel weights predicting SM, and results of the between-region tests of ‘boost’ to model evidence for PFC plus PVC models compared to PVC-only. See text for details. Alpha = .0125. SD = standard deviation. SM = subsequent memory. n=119.

#### Testing posterior-anterior shift

The foregoing analyses provide strong evidence that the increase in (univariate) PFC activity observed in this task did not reflect compensation. Nonetheless, the PASA theory is more general, describing a shift in relative reliance on posterior and anterior regions with age, which need not be compensatory, but could reflect differential age effects in posterior and anterior cortices. In other words, the relative involvement of anterior versus posterior regions could increase with age, even if the absolute involvement of both decreased with age. Direct comparison of the mean univariate activation between ROIs did not reveal any evidence for such relative differences in age effects, with strong Bayesian evidence against the predicted greater age-related increase in PFC (BF_01_ for null hypothesis = 25; Table 2). We next tested for a shift in the relative multivariate information between regions. In the separate MVB models, the age-related reduction in spread of weights was numerically greater in PFC than PVC (linear age effect on PFC-PVC difference, *t*(118) = 4.16, p = .0437); see Table 2. We also measured the proportion of top-weighted voxels (> 2 standard deviations above the mean) that were located in PFC as opposed to PVC in the joint PVC-PFC model. This proportion decreased significantly with age (overall model, *F*(2,116) = 3.27, p = .0415; linear *t*(118) = -2.55, *r*^2^adj = .359, p = .0119; quadratic *t*(118) = -.106, p = .915), with mean 52.4% of top voxels located in PFC in the younger tertile (SD = 9.09; 18-45 years) and 47.1% in the older tertile (SD = 8.57; 65-88 years), although this was no longer significant when item misses were excluded (overall model, *F*(2,105) = 2.60, p = .0799, linear *t*(107) = -1.86, p = .0638). Thus, there was no evidence that in older age there is a general shift in the areas contributing to subsequent memory from posterior to anterior (though see Experiment 2 below).

### Experiment 2: Short-term Memory (STM) Maintenance

#### Behavioral results

For the visual STM task, analysis of the effects of increasing load on performance showed a strong age-related increase in the effect of load on accuracy, measured using the root mean squared error of the estimated dot direction relative to the actual dot direction in degrees (Fig 1b). As expected, older people showed a larger increase in error at load = 3 compared to load = 1 (linear contrast) than younger people (for linear age-by-load interaction, *t*(95) = 5.53, p < .0001, *r*^2^adj = .192; quadratic *t*(95) = =1.27, p = .203), although some age-related decrement in accuracy was present even at load = 1 (for linear effect of age, *t*(95) = 2.607, p = 0110, *r*^2^adj = .0382; quadratic *t*(95) = 0.388, p = .699).

#### Testing compensation

For STM, standard univariate activation analyses showed that increasing load elicited activity increases during the maintenance period which varied according to age in both ROIs. As in the LTM experiment, and consistent with the PASA account, PFC activation increased with age, particularly in later years (linear *t*(95) = 3.01, p = .003; quadratic *t*(95) = -0.505, p = .615) (Fig 2c, Table 4). Unlike for LTM encoding, there was also a significant increase in load-related PVC activation over the lifespan (linear *t*(95) = 4.28, p < .0001; quadratic *t*(93) = -0.988, p = .324; see Table 4.

**Table 4.** Age effects on mean univariate SM effects and spread of multivariate SM effects in the STM task. PFC-PVC refers to analyses where the dependent variable was the difference in each measure between PFC and PVC. SD = standard deviation. n = 96. R^2^adj = the unbiased estimate of the amount of variance explained in the population.

| ROI/<br>measure | Model |  | Linear |  |  | Quadratic |  |  |
| --- | --- | --- | --- | --- | --- | --- | --- | --- |
| | *F* | *p* | *t* | $r^2_{adj}$ | *p* | *T* | $r^2_{adj}$ | *p* |
| Mean univariate STM activation |  |  |  |  |  |  |  |  |
| PFC | 4.57 | .0128 | 3.01 | .0553 | .00336 | -0.505 | -- | .615 |
| PVC | 9.43 | .000187 | 4.28 | .119 | < .0001 | -0.988 | -- | .324 |
| PFC-PVC | .606 | .548 | -0.587 | -- | .559 | -0.878 | -- | .380 |
| Multivariate spread (SD) of STM activity |  |  |  |  |  |  |  |  |
| PFC | 13.4 | < .0001 | .662 | -- | .507 | -5.03 | .162 | < .0001 |
| PVC | 9.30 | .000210 | -1.01 | -- | .308 | -4.07 | .108 | < .0001 |
| PFC-PFC | 3.00 | .0547 | .674 | -- | .497 | 2.26 | .0250 | .0244 |

Separate MVB analysis in each ROI showed a similar pattern of age effects to the LTM task. In both PFC and PFC, the spread (SD) of individual voxel weights predicting increased STM load was particularly reduced during later life, with a significant quadratic component (PVC: linear *t*(95) = -1.01, p = .308; quadratic *t*(95) = -4.07, p < .0001; PFC: linear *t*(95) = 0.662, p = .507; quadratic *t*(95) = -5.03, p < .0001) (see Fig 2d and Table 4). The result for PVC was unchanged by removing a subset of subjects with very low values (i.e., SD weights < .0005; for quadratic age effect *t*(69) = -5.32, p = .0012). As found for LTM encoding, the direction of the effect in PFC was contrary to a compensatory PASA shift, i.e., PFC voxels contributed less to the cognitive task in old age. Again, the spread of univariate effects did not show the same effects of age (in PVC linear *t*(95) = 1.52, p = .134; quadratic *t*(95) = 0.831, p = .406; in PFC, linear *t*(95) = -0.471, p = .641; quadratic *t*(95) = 1.70, p = .0912).

As for LTM encoding, we used model comparison of a joint PVC-PFC model with a PVC-only model to directly evaluate the compensatory PASA hypothesis. The results were similar to the LTM experiment: The “boost” to prediction of the cognitive variable obtained by adding PFC to the model showed a significant age-related *reduction* in the probability of a boost for STM load (in an ordinal regression, *t*(95) = -2.00, p = .0479). The Bayes Factor provided strong evidence against the compensatory hypothesis of an increased boost (for unidirectional hypothesis, BF = 10.2), although evidence was only anecdotal for the presence of an age-related reduction in boost (for bidirectional hypothesis, BF = 1.81). Thus, like for the LTM experiment, the<u>r</u>e was clear evidence against a compensatory increase in prefrontal contribution to the task with age.

#### Testing sub-regions within PFC

Again, we examined the four prefrontal subregions separately to assess whether the findings were driven by a specific part or parts of the large ROI (Table 5). For this experiment, the age-related increase in univariate activation was not separately significant in any subregion, which may have reflected relatively distributed effects and the more inclusive selection of ‘active’ voxels. As for the LTM experiment however, overall age effects on the spread of multivariate voxel weights were significant in three subregions, and those in IFG were similar in magnitude and form, suggesting reductions in spread across PFC, with no major between-subregion differences. Moreover, all ROIs showed Bayes Factors of at least 6 against a boost to model evidence from adding PFC to the posterior-only models predicting increasing STM load, again consistent with the overall results.

**Table 5.** Age effects for PFC subregions in the visual short-term memory task. The table lists mean univariate activation during maintenance in response to increasing VSTM load, the spread (SD) of MVB voxel weights predicting linearly increasing VSTM load, and results of the between-region tests of ‘boost’ to model evidence for PFC plus PVC models compared to PVC-only. See text for details. Alpha = .0125. SD = standard deviation. n=96.

| ROI/ measure | Model |  | Linear |  |  | Quadratic |  |  |
| --- | --- | --- | --- | --- | --- | --- | --- | --- |
| | $F$ | $P$ | $t$ | $r^2_{adj}$ | $p$ | $t$ | $r^2_{adj}$ | $p$ |
| **BA10** |  |  |  |  |  |  |  |  |
| Mean univariate | 1.90 | .155 | 1.79 | -- | .0787 | -0.937 | -- | .352 |
| MVB spread (SD) | 12.7 | < .0001 | -1.41 | -- | .158 | -4.69 | .171 | < .0001 |
| PFC boost $t$ | -- | -- | -1.48 | -- | .142 | -1.00 | -- | .320 |
| PFC boost $BF_{01}$ | -- | -- | 10.0 | -- | -- | -- | -- | -- |
| **IFG** |  |  |  |  |  |  |  |  |
| Mean univariate | 2.64 | .0767 | 1.99 | -- | .0512 | 0.997 | -- | .323 |
| MVB spread (SD) | 6.26 | .00282 | -0.787 | -- | .427 | -3.35 | .0864 | .00109 |
| PFC boost $t$ | -- | -- | -1.11 | -- | .270 | -1.10 | -- | .274 |
| PFC boost $BF_{01}$ | -- | -- | 7.69 | -- | -- | -- | -- | -- |

**Table 5.** Age effects for PFC subregions in the visual short-term memory task.
| **MFG** |  |  |  |  |  |  |  |  |
| --- | --- | --- | --- | --- | --- | --- | --- | --- |
| Mean univariate | 2.08 | .131 | 2.03 | -- | .0459 | -0.447 | -- | .654 |
| MVB spread (SD) | 11.0 | < .0001 | 0.300 | -- | .762 | -4.60 | .165 | < .0001 |
| PFC boost $t$ | -- | -- | -1.38 | -- | .171 | -0.788 | -- | .433 |
| PFC boost $BF_{01}$ | -- | -- | 6.25 | -- | -- | -- | -- | -- |
| **SFG** |  |  |  |  |  |  |  |  |
| Mean univariate | 2.64 | .0770 | 2.27 | -- | .0264 | 0.241 | -- | .812 |
| MVB spread (SD) | 7.37 | .00106 | -1.57 | -- | .116 | -3.34 | .0858 | .00111 |
| PFC boost $t$ | -- | -- | -2.73 | .0528 | .00755 | -0.720 | -- | .473 |
| PFC boost $BF_{01}$ | -- | -- | 14.3 | -- | -- | -- | -- | -- |

#### Testing posterior-anterior shift

Finally, even if age increased activity and decreased multivariate information in both PFC and PVC, it is possible that the PFC:PVC ratio of activity and/or multivariate information increases with age, consistent with the general PASA claim. As for LTM encoding, there was no evidence that age effects on (univariate) activation in the two ROIs differed, i.e. the increase in activation in older people was similar in magnitude (Table 4; BF_01_ for null hypothesis over prediction of a greater age-related increase in PFC = 33.3). However, multivariate analysis revealed a picture different from that in the LTM task. In the separate MVB models, the age-related reduction in spread of weights showed a stronger quadratic component in PFC than PVC (for PFC-PFC, quadratic *t*(95) = 2.26, p = .0244; Table 4). More clearly, when examining the location of top-weighted voxels from the joint PFC+PVC model, a higher proportion were located in PFC in older age (for model, *F*(2,93) = 22.4, p < .0001, linear *t*(95) = 3.72, p < .001, quadratic *t*(95) = 5.20, p < .0001), with mean 69.1% of top voxels located in PFC in the younger tertile (SD = 14.1; 25-43 years) but 81.0% in the older tertile (SD = 13.3; 66-85 years). Thus while the STM experiment, like the LTM experiment, found decreases in absolute PFC (and PVC) involvement in old age, the relative involvement of PFC versus PVC voxels (at least in terms of those with high weights in the joint model) did increase with age, unlike in the LTM experiment. This provides some support for a PASA pattern in this task, even though there was no evidence that this greater PFC involvement was compensatory.

## Discussion

This study investigated the proposal that there is a posterior-to-anterior shift in task-related brain activity during aging, with the greater reliance on prefrontal cortex in older age reflecting compensatory mechanisms. We tested the predictions of this PASA theory with data from two memory tasks that were conducted on independent and relatively large population-derived adult lifespan samples. Using novel model-based multivariate analyses, we provide direct evidence against a compensatory posterior-to-anterior shift. Instead, our data suggest that the increased prefrontal activation reflects less specific or less efficient activity, rather than compensation.

The results of our standard univariate activation analyses are consistent with previous studies showing age-related increases in activation in prefrontal cortex, which form the basis of the PASA theory (Grady et al., 1994; Davis et al., 2008). Many studies have found such increases in PFC activation in different cognitive tasks, although regional reductions in activation are also found (e.g. see Rajah and D’Esposito, 2005; Spreng et al., 2010; Li et al., 2015). We found such age-related increases in both the PFC activation associated with trials that were later remembered many minutes later (in the LTM experiment) and the activation associated with maintaining increasing numbers of items for a few seconds (in the STM experiment). We also further generalized previous findings across PFC sub-regions, in that the increased activation was reliable across lateral, anterior and superior prefrontal areas (although in the LTM experiment, it was mainly driven by inferolateral and anterior PFC).

Importantly, despite this increased univariate activity, multivariate analysis of both experiments showed that with increasing age, PFC possessed less, rather than more, information about the cognitive outcome. This reduced pattern information was evident both in terms of the spread of voxel weights (Figure 2) and the lack of a meaningful boost to model evidence when adding PFC voxels to the model (Figure 3). The latter type of inference was made possible by our novel use of multivariate Bayesian (MVB) classification.

If the increased prefrontal activation with age is not compensatory, then what does it reflect? One possibility is that neural function is less efficient, such that a greater BOLD signal is required for the same level of computation, i.e, less “bang for the buck” for the same level of neural activity (Grady, 2008; see also Rypma and Esposito, 2000; Morcom et al., 2007; Reuter-Lorenz and Campbell, 2008; Nyberg et al., 2014). This could reflect the proposed greater sensitivity of prefrontal cortices than other brain regions to aging (West, 1996; Glisky et al., 2001; Raz and Rodrigue, 2006). Another possibility is that PFC activity becomes less specific with age, as might be expected by theories of age-related dedifferentiation, particularly in complex cognitive functions (Li et al., 2001; Park et al., 2004; Carp et al., 2011; Abdulrahman et al., 2014). Partial support for the latter comes from the LTM experiment, where the negative effect of age on the spread of multivariate weights across voxels was accompanied by a positive effect of age on the spread (as well as mean) of univariate activity. This suggests that, while more voxels showed substantial (positive or negative) activity related to subsequent memory in older people, these additional responses were redundant, with fewer voxels contributing uniquely to memory encoding, as expected if the increased prefrontal activity is less specific. In the STM experiment, on the other hand, the spread of univariate responses was age-invariant, suggesting a more spatially uniform increase in response to load with age, although the MVB results suggested that – just as in the LTM task – this increased response carried less information. Whether the present results reflect reductions in efficiency or reductions in specificity, they are more consistent with the general idea of brain maintenance (Nyberg et al., 2014) – that cognitive function in older age is determined by the ability to maintain a youth-like brain – than with the idea associated with PASA of functional compensation by anterior brain regions.

Despite age-related decreases in overall multivariate information in both PFC and PVC, it is possible that the *relative* contribution of anterior regions to cognitive tasks could increase with age. There is some evidence for such a shift from studies showing crossover effects in which age-related decreases in posterior cortical activity occur alongside age-related increases in PFC (e.g., Grady et al., 1994; Davis et al., 2008; see also recent meta-analysis by Maillet and Rajah, 2014). However, our univariate activation analyses showed little evidence of such a relative posterior-to-anterior shift: despite increased prefrontal activation, age effects on univariate activation in PFC and PVC did not differ significantly in either experiment. In terms of multivariate information, the LTM experiment actually showed, if anything, a decrease rather than increase in the contribution of PFC relative to PVC. The only comparison that provided some support for a relative increase in anterior contribution was for multivariate information about load in the STM experiment. Thus the direction of any *relative* shift in reliance on PFC versus PVC with age seems to be task-dependent, as opposed to the task-general posterior-to-anterior shift claimed by PASA (Davis et al., 2008; see also Ford and Kensinger, 2017). This is consistent with other meta-analyses, which have found age-related decreases as well as increases in activation, depending on the task (Spreng, Wojtowicz, & Grady, 2010; Li et al., 2015). Moreover, most studies have not made the direct statistical comparisons needed to test for anterior-posterior differences in the absence of crossover effects (see Morcom & Johnson, 2015). A strength of our approach is that our analyses encompassed large ROIs in both anterior and posterior cortices, as well as direct comparisons between the two.

In summary, our data replicate an increase in PFC activity over the adult lifespan, but do not support the idea that this reflects a compensatory posterior-to-anterior shift, at least in the context of the two memory tasks considered here. Our results are inconsistent both with the proposal that the increased activity is compensatory, and with a generalized shift with age to greater relative reliance on prefrontal cortex. The data are most parsimoniously explained by reduced efficiency or specificity of neural responses, reflecting primary age-related deleterious changes in posterior as well as prefrontal cortex which vary in their relative magnitudes according to the task. Our results therefore help to adjudicate between competing accounts of neurocognitive aging, while also illustrating the more general ability of MVB to compare models that comprise different sets of voxels, thereby offering an exciting new general way to test the relative contributions of brain regions to cognitive outcomes.

## Acknowledgments

A.M.M. is a member of the University of Edinburgh Centre for Cognitive Ageing and Cognitive Epidemiology (CCACE), part of the UK cross-council Lifelong Health and Wellbeing Initiative, Grant number G0700704/84698. The Cambridge Centre for Ageing and Neuroscience (Cam-CAN) research was supported by the Biotechnology and Biological Sciences Research Council (grant number BB/H008217/1). The full Cam-CAN author list can be found here: http://www.cam-can.org/index.php?content=corpauth. R.N.A.H. is funded by the Medical Research Council (SUAG/010 RG91365) with additional support by the European Union’s Horizon 2020 research and innovation programme (grant agreement No 732592).

## Author contributions

A.M.M. designed research; A.M.M., R.N.A.H. and Cam-CAN performed research; A.M.M. and R.N.A.H. analyzed data; A.M.M. and R.N.A.H. wrote the paper.

